# Techniques for Developing Reliable Machine Learning Classifiers Applied to Understanding and Predicting Protein:Protein Interaction Hot Spots

**DOI:** 10.1101/2022.12.26.521948

**Authors:** Jiaxing Chen, Leslie A. Kuhn, Sebastian Raschka

## Abstract

With machine learning now transforming the sciences, successful prediction of biological structure or activity is mainly limited by the extent and quality of data available for training, the astute choice of features for prediction, and thorough assessment of the robustness of prediction on a variety of new cases. Here we address these issues while developing and sharing protocols to build a robust dataset and rigorously compare several predictive classifiers using the opensource Python machine learning library, scikit-learn. We show how to evaluate whether enough data has been used for training and whether the classifier has been overfit to training data. The most telling experiment is 500-fold repartitioning of the training and test sets, followed by prediction, which gives a good indication of whether a classifier performs consistently well on different datasets. An intuitive method is used to quantify which features are most important for correct prediction.

The resulting well-trained classifier, *hotspotter*, can robustly predict the small subset of amino acid residues on the surface of a protein that are energetically most important for binding a protein partner: the interaction hot spots. *Hotspotter* has been trained and tested here on a curated dataset assembled from 1,046 non-redundant alanine scanning mutation sites with experimentally measured change in binding free energy values from 97 different protein complexes; this dataset is available to download. The accessible surface area of the wild-type residue at a given site and its degree of evolutionary conservation proved the most important features to identify hot spots. A variant classifier was trained and validated for proteins where only the amino acid sequence is available, augmented by secondary structure assignment. This version of *hotspotter* requiring fewer features is almost as robust as the structure-based classifier. Application to the ACE2 receptor, which mediates COVID-19 virus entry into human cells, identified the critical hot spot triad of ACE2 residues at the center of the small interface with the CoV-2 spike protein. *Hotspotter* results can be used to guide the strategic design of protein interfaces and ligands and also to identify likely interfacial residues for protein:protein docking.

## Introduction

### Characteristics of protein:protein interfaces

Protein-protein interactions *in vivo* are fundamental for coordinating and regulating biological pathways, including the COVID-19 coronavirus spike protein interaction with the human ACE2 receptor, which mediates infection [1, 2]. Protein-protein interfaces [3] on average bury 1200– 2000 Å^2^ of the surface [4], involving an average of 87 atoms from 22 residues per protein [5]. Although protein interfaces can be large, only a few residues – on average, 9.5% of the interfacial residues – contribute the majority of binding affinity [6] and are referred to as hot spots of interaction. Mutation of hot spot residues to alanine often weakens the binding free energy between protein partners by 2.0 kcal/mol or more [7]. Because of their significant stabilizing effect on protein-protein interactions, identification of hot spot residues is essential for protein engineering and drug discovery [8, 9]. In fact, drug-like small molecules usually bind to hot spots on protein interfaces [8], and mutation of hot spot residues can either reduce or strengthen protein-protein interactions.

### Approaches for predicting protein:protein interaction hot spots

Experimental methods to systematically identify interaction hot spots are laborious and expensive, making them inaccessible to many research groups [6]. Alanine scanning mutagenesis involves plasmid construction, protein expression and purification, followed by *in vitro* assays to measure binding affinity [6] for each mutation in a protein. The experimental challenges have motivated the development of computational methods to pinpoint the most likely hot spot residues for more targeted experiments. Development of hot spot predictors can, in turn, clarify the features dominating the free energy of binding between proteins.

These predictive methods fall into three categories: molecular simulations, empirical or physical models of interaction, and machine learning methods that learn from the features of known hot spots. Molecular dynamics can simulate structural fluctuations and transitions in the wild-type versus alanine mutant structures, allowing an estimate of the change in protein binding free energy from a mutation. However, this approach is computationally intensive and does not provide information about hot spot determinants in general [10, 11]. Knowledge-based and physical models make use of energy terms with optimized weights fit to experimental binding data [12, 13]. The constructed energy functions are then applied to calculate the change in free binding energy between a wild-type residue and its alanine mutation in a given protein [14]. This approach dramatically reduces the computational time relative to molecular dynamics; however, the performance in distinguishing hot spots from non-hot spots is variable.

Recently, a variety of machine learning methods including classifiers have been used to predict hot spots, such as decision trees [13], support vector machines [11, 15–17], ensemble methods [11], and extreme gradient boosting methods [18]. Classifiers are commonly used predictors that seek to divide cases into categories, e.g., hot spot or non-hot spot residues. Machine learning models are trained on a dataset with labeled hot spot residues and non-hot spot residues. Feature values based on sequence, structure, chemistry, and/or interaction energy are provided by the user to characterize each site. Once calculated, these feature values are used as inputs for the classification of hot spot and non-hot spot residues. Compared to molecular simulations and physical models, machine learning methods are generally more efficient and have better predictive performance. Some machine learning approaches provide information about the most important features for prediction, whereas others provide reliable predictions without human-interpretable models.

### Dataset size and redundancy in training and test sets for machine learning

The size of the dataset used for training and unbiased testing of a machine learning method is important for its performance on new data. Of particular importance is whether the data represents an even sampling of the diversity of cases on which the predictor will be applied. A model constructed using a small dataset and a large number of features has a high risk of overfitting. Overfitting occurs when the predictive model uses the large number of features during training to closely fit the cases in the relatively small training dataset. This results in effectively memorizing details of specific cases rather than discerning underlying trends in the data, leading to much worse performance on new data [19]. Outliers also have a large impact on a predictive model when using a small dataset and hence can skew the results, whereas they are much less of an issue for models trained on large datasets. Use of a small training or test dataset similarly does not allow a clear appraisal of predictive quality in general.

Homology between protein sites also needs to be addressed carefully to avoid predictive bias when constructing datasets for training and testing. The issue is more complicated when considering interaction sites, which include contacts between *two* proteins. Redundancy of sites within a training or test set will overweigh that type of site in the results, skewing the performance metrics. Including redundant sites between the test set and the training set can easily occur when one database of binding sites is used for training and another is used for prediction, without careful comparison and culling of the same sites between the two sets. Predictors can learn the features of the input case during optimization, making the same or very similar case in the test set easy to predict. This leads to a false boost in performance relative to what will be found for actual new cases. The same bias can also occur in binding site prediction methods that simply use 35% or higher sequence identity between two proteins (or a similar rule) as the criterion for removing redundant sites. This criterion is inadequate because protein binding site residues are often highly conserved even when two proteins are less than 35% identical in overall sequence.

The above considerations indicate that identity within and between test and training cases during predictor development needs to be considered at the site level: a single amino acid and its contacts with the other protein in the complex. Just as redundant sites are possible when overall sequence identity is used to assess non-identity between sites, it is also possible to accidentally exclude unique sites. Typically, many different residues are probed in a single protein to measure changes in binding free energy and identify hot spots. We would not want to exclude an entire identical protein chain between data sets if different residues were probed by mutation. Thus, a focus on the residue and residue-contact level rather than chain level is required to assess redundancy.

Another issue to consider when comparing sites is that residues in homologous or identical proteins are often numbered differently between structure files in the Protein Data Bank (PDB; rcsb.org; [20]). Different residue numbering often occurs between mutation databases and sequence or PDB files of the same protein, too, because different length protein constructs and numbering schemes are used by different research groups. Mis-indexing sites between proteins is more of a problem when sites in the same protein from multiple databases are assembled into a larger dataset. As a result, building a complete but non-redundant dataset involves careful curation of the residue entries, site by site. Aside from small database size, a frequent weakness in publications on hot spot prediction is that the level of thoroughness in removing redundant entries in training and test sets is unclear.

### Databases of experimental data for hot spot training and testing

Most machine learning methods for predicting protein hot spots have been trained on data from one or more of these binding energy databases: Alanine Scanning Energetics Database (ASEdb) [7], the Binding Interface Database (BID) [21], and the Structural Kinetic and Energetic Database of Mutant Protein Interactions (SKEMPI) [22]. Our work incorporates and culls redundant examples of data from these sets as well as including the Antibody Binding Mutational Database (AB) [23]. Antibody:protein complexes differ in structure from other complexes due to binding the antigen protein via six loops on the antibody rather than forming a relatively flat interface. Antibody complexes are an especially important class of interaction to understand for vaccine and biologic drug development [24].

We address the above considerations during training, testing and comparing the results of five prominent machine learning classifier methods for predicting interaction hot spots from 1,046 non-redundant residues with experimental binding free energy data for comparison. In the process, we show how to evaluate if there is sufficient training data; whether overfitting has occurred due to insufficient training data or too many trainable features or parameters; which features are the most important for prediction; and how consistently and well a predictor performs on new data. We also discuss how to choose performance measures (metrics) which are the most appropriate for problems like this, in which the true positives (hot spots) are buried in a large number of other cases (non-hot spots). As we’ll discuss, underprediction of rare true positive cases and overprediction of false positives are serious problems hidden by the performance metric most commonly used in hot spot publications: accuracy (the percentage of all cases that are correctly classified).

## Materials and Methods

### Constructing training and testing datasets and removing redundant sites

Single-site alanine mutation data were collected and pooled from the Alanine Scanning Energetics Database (ASEdb) [7], the Structural Kinetic and Energetic Database of Mutant Protein Interactions (SKEMPI) [22], the Antibody Binding Mutational Database (AB) [23], and the Binding Interface Database (BID) [21]. Following careful indexing of residue numbers for the same protein in the different datasets including the PDB, if a site (defined as the same residue position in the same protein based on sequence or structural comparison) appeared in more than one of the mutational databases, it was only included once in the pooled training and test sets. If the same residue in the same protein occurred in complex with a different protein, resulting in different contacts for the residue and a separately measured ΔΔG_binding_ for the protein complex, it was kept. Sites without crystal structures (since structural features are used in one version of *hotspotter*) or which occurred in proteins with multiple mutations in a single binding free energy experiment (such that the contribution of each single residue to the measured ΔΔG_binding_ could not be defined) were removed. (Note 1) In the SKEMPI database, the affinity value (K_d_) of the wild-type complex and the affinity value of the mutant complex are given rather than the change in binding free energy upon mutation. For the SKEMPI cases, the following equations were used to obtain ΔΔG values from the K_d_ values, with room temperature (298 K) used for calculation:

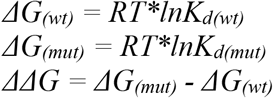

As a result, from 97 protein complexes (Table 1) we obtained 1,046 sites at which alanine has been substituted for the wild-type residue, along with the experimental measurement of the mutation-caused change in ΔG_binding_ between the two proteins, referred to as ΔΔG_binding_. Residues mutated to alanine with ΔΔG_binding_ greater than or equal to 2.0 kcal/mol, reflecting decreased affinity between the proteins upon mutation, were defined as hot spots. Those with values between 1.0 and 2.0 kcal/mol were labelled as weak hot spots, and those with ΔΔG less than 1.0 kcal/mol were labelled as non-hot spots. Mutations in the BID database are annotated as “strong”, “intermediate”, “weak”, “insignificant”, “negative”, “negative-strong”, “negative-intermediate”, or “negative-weak” rather than by numeric values. Residues labeled as “strong” were defined as hot spots and those labeled as “insignificant”, “negative”, “negative-strong”, “negative-intermediate”, or “negative-weak” were defined as non-hot spots.

**Table 1.**
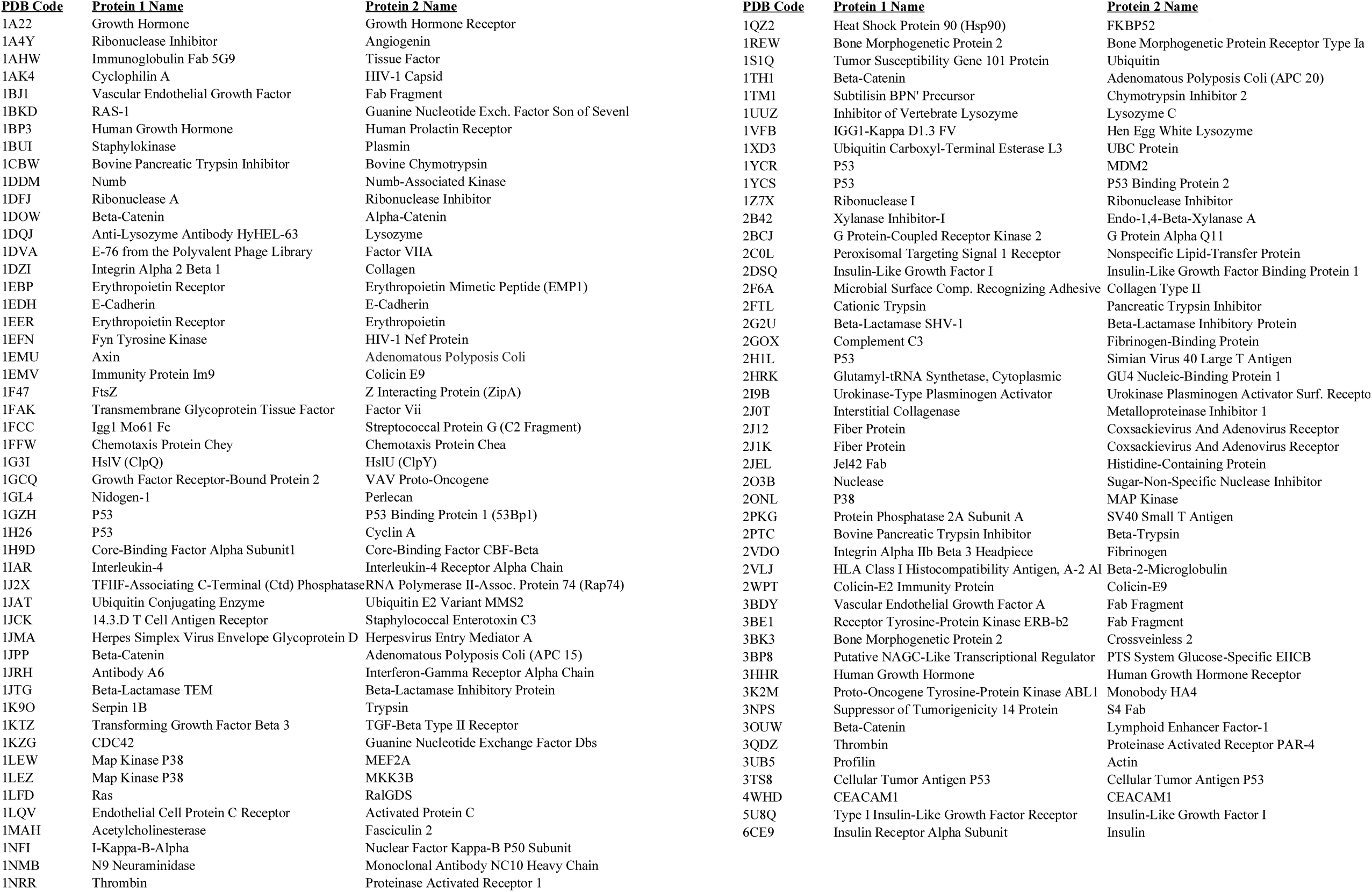
97 protein:protein complexes used to create a feature dataset of 1.046 protein hot spot and non-hot spot sites.

After initial testing of several classifiers, the weak hot spot cases were merged into the hot spot category, resulting in better discrimination than grouping them with non-hot spots. The final 1,046 sites consisted of 449 hot spots and 597 non-hot spots. This does not reflect the natural prevalence of hot spots relative to non hot spots in the 97 proteins. The goal was to build the largest dataset of non-redundant hot spots possible (the limiting, important case) and then construct a similar number of non-redundant non-hot spots for balance. This allows the classifier to learn the determinants of both classes. The data were split into training and test sets randomly, with 70% of the hot spot sites used for training and 30% for testing. A 70%:30% split between training and test sets was also used for the non-hot spot sites.

### Characterizing the features of each site as the basis for prediction

Since mutation of hot spot residues can cause a significant increase (weakening) in binding free energy of complex formation between two proteins, we hypothesized that hot spot residues form more non-covalent interactions than other surface or interfacial residues. We used ProFlex v. 5.1, software created by our lab [25, 26] and available at https://github.com/psa-lab/ProFlex, to calculate three bond and contact features for each residue: average bond number (the average number of covalent, hydrophobic, and hydrogen-bond contacts for atoms in the residue), the number of hydrogen bonds formed (considering the detailed geometry and length of bonds involving the donor, hydrogen and acceptor atoms), and the number of interatomic hydrophobic contacts, referred to as hydrophobic tethers in ProFlex. One ProFlex feature that was calculated, average bond weight, was not kept in our classifiers because it did not enhance the quality of prediction. To determine the energy level for use in ProFlex for calculating features, the Hether script distributed with ProFlex was used to identify the energy at which the number of rigid clusters of protein atoms changed most.

For the fourth feature, crystallographic temperature factor values, also known as B factors or B values, which experimentally quantify the mobility of each atom in the crystal structure, were extracted from the PDB file. The B values were used to calculate B’ for the side chain, in which the B values of the atoms in each side chain are averaged and divided by the average B value of all side chains in the protein, and then normalized by the average side-chain occupancy values for the same atoms. The occupancy data is also provided in PDB files [27]. This B’ normalization scales mobility values so they are on a consistent scale in different PDB entries [27, 28].

Some amino acid types, such as tryptophan, arginine, and tyrosine, are reported to be enhanced in known protein:protein hot spots [6, 29]. To test this hot spot propensity as a fifth feature, we measured the hot-spot probability of each residue. This was calculated for each amino acid type as the percentage of hot spot occurrences for that amino acid in in all the non-redundant mutations from ASEdb, AB, BID, and SKEMPI databases. Later, this feature was replaced by the residue type (e.g., Ala, Arg, etc.), encoded as a one-hot bit string (10000… for Ala, 01000… for Arg, etc.), to avoid any prior-knowledge bias and allow the machine learning method to discern the relative importance of residue type for prediction.

The sixth feature was based on the observation that residues involved in protein or ligand binding tend to be highly conserved in amino acid type across evolution. To measure residue conservation, we used the online ConSurf server (https://consurf.tau.ac.il; [30]) in default mode to generate a multiple sequence alignment of diverse homologs for each of the 97 PDB files in our dataset and assign a conservation value to each residue (from 1=most variable in amino acid type in the multiple sequence alignment to 9=most conserved).

The final two residue features, accessible surface area (ASA) in the protein:protein complex and secondary structure type were calculated for the wild-type residue in each site with GROMACS version 2021.2. The four-state secondary structure type (T for turn, H for helix, S for strand, and – for other) was computed using the DSSP option in GROMACS [31–33], and the solvent accessible surface area was calculated using SASA in GROMACS [34]. All features used to characterize sites for hot spot prediction are intuitive structural or sequence-based features, which is beneficial for their experimental interpretation and use in guiding protein design.

Two spreadsheets (one for the training dataset, the other for testing), including the 1,046 sites with their feature values and experimental ΔΔG_binding_ or binding affinity values from the original database (SKEMPI, ASEdb, BID, or AB), are provided on GitHub (https://github.com/Raschka-research-group/hotspotter/tree/main/experiments/dataset). Additionally, the spreadsheets include three columns used in early experiments that were not used in recent or final classifiers; average bond weight, hot spot ratio, and a 2-class value arising from experimental data (value of 1 for strong hot spots with experimental ΔΔG_binding_ ≥≥ 2 kcal/mol and value of 0 for non-hot spot residues with ΔΔG_binding_ ≤ 0 kcal/mol, with the intermediate weak hot spots assigned blank values). The 3-class column (value of 0 for non-hot spots, value of 1 for weak hot spots with experimental ΔΔG_binding_ values between 0 and 1, and value of 2 for strong hot spots with ΔΔG_binding_ ≥ 2 kcal/mol) was used to define the hot spot class of each site when training the current classifiers and to assess the correctness of test case predictions as hot spot (including both weak and strong hot spots) or non-hot spot. The overall pipeline for this hot spot prediction approach is illustrated in Figure 1.

**Figure 1.**
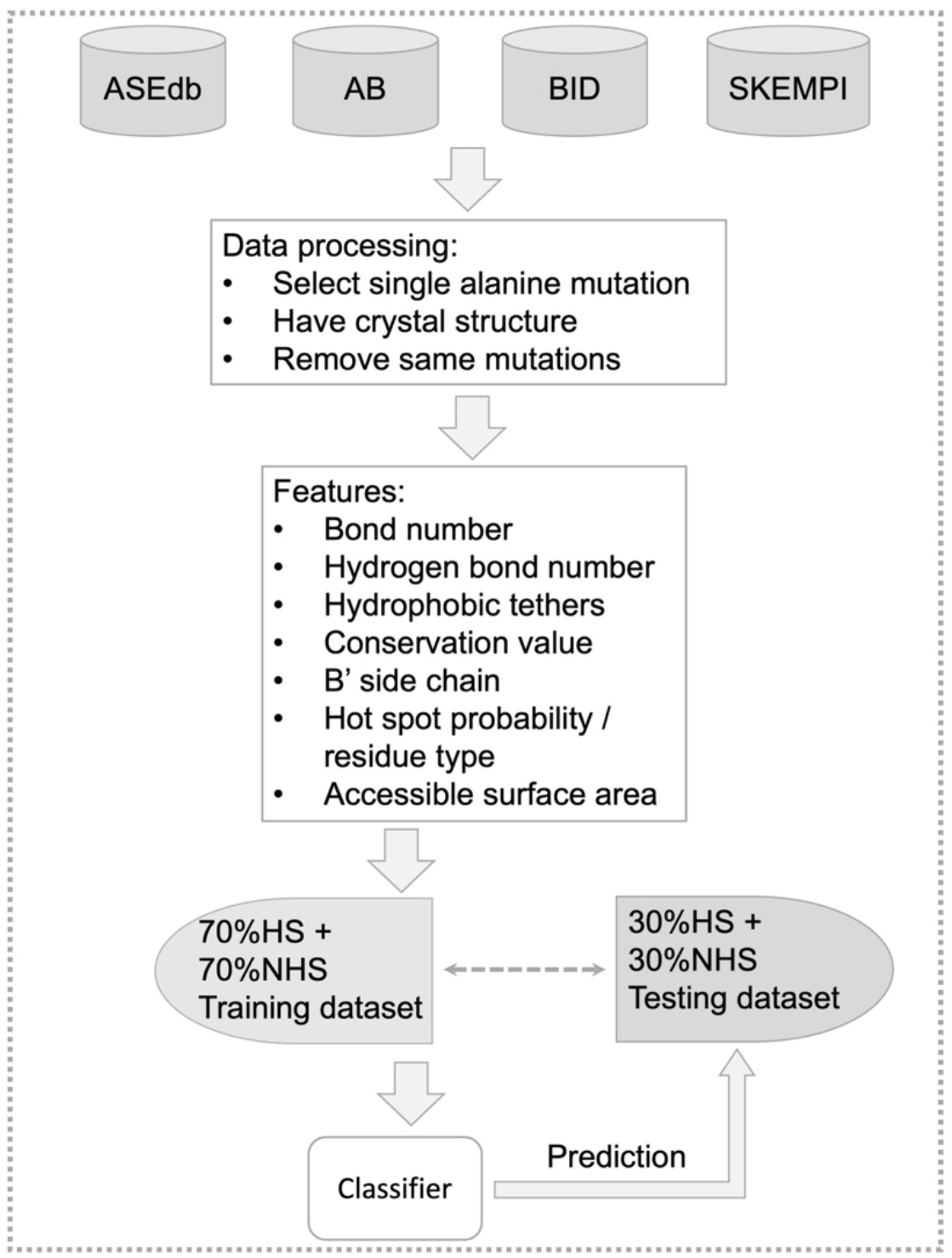
Training and testing of *hotspotter* on the features of 1,046 mutation sites in 97 protein:protein complexes.

### Evaluating a panel of machine learning methods for hot spot prediction

Five different machine learning classifiers, logistic regression, random forest, gradient boosting, support vector machine, and multilayer perceptron [35], were evaluated for training and testing, given the feature values characterizing the 1,046 sites. Logistic regression is a binary classifier using a sigmoidal activation function to best fit a binary output (e.g., hot spot or non-hot spot), based on the feature values and known classes of the training set. The random forest algorithm assembles a set of decision trees trained on the feature values and known classes (hot spot or non-hot spot) of random bootstrap samples of the training set. For classifying each example in the test set, the random forest method determines the binary class label via a majority vote: each tree in the random forest ensemble makes a prediction, and the most frequent class label among all trees is returned as the predicted label. Random forests are less prone than single decision trees are to overfitting to the training data.

Gradient boosting is another ensemble technique based on decision trees. Unlike random forest, an ensemble method that trains very deep decision trees independently, gradient boosting trains many short trees sequentially. The first tree in this sequence is trained to predict the class labels. Then the subsequent trees are trained on the errors of the previous trees in an iterative fashion. The fourth method used was a support vector machine, which maps training examples to points in space based on their feature values in such a way that the width of gap between samples in the two categories is maximized (e.g., hot spot versus non-hot spot). Test examples are then mapped into the same space, and their categories are predicted based on which side of the gap they occur. The fifth classifier tested was a multilayer perceptron, a type of feed-forward, fully connected, multilayer neural network. To train and evaluate the machine learning algorithms mentioned above, as described in the steps in Figure 2, we used scikit-learn version 1.1.2 (https://scikit-learn.org; [36]). All experiments relating to training, testing, and evaluating the results of classification methods are provided as code on GitHub (https://github.com/Raschka-research-group/hotspotter/tree/main/experiments).

**Figure 2.**
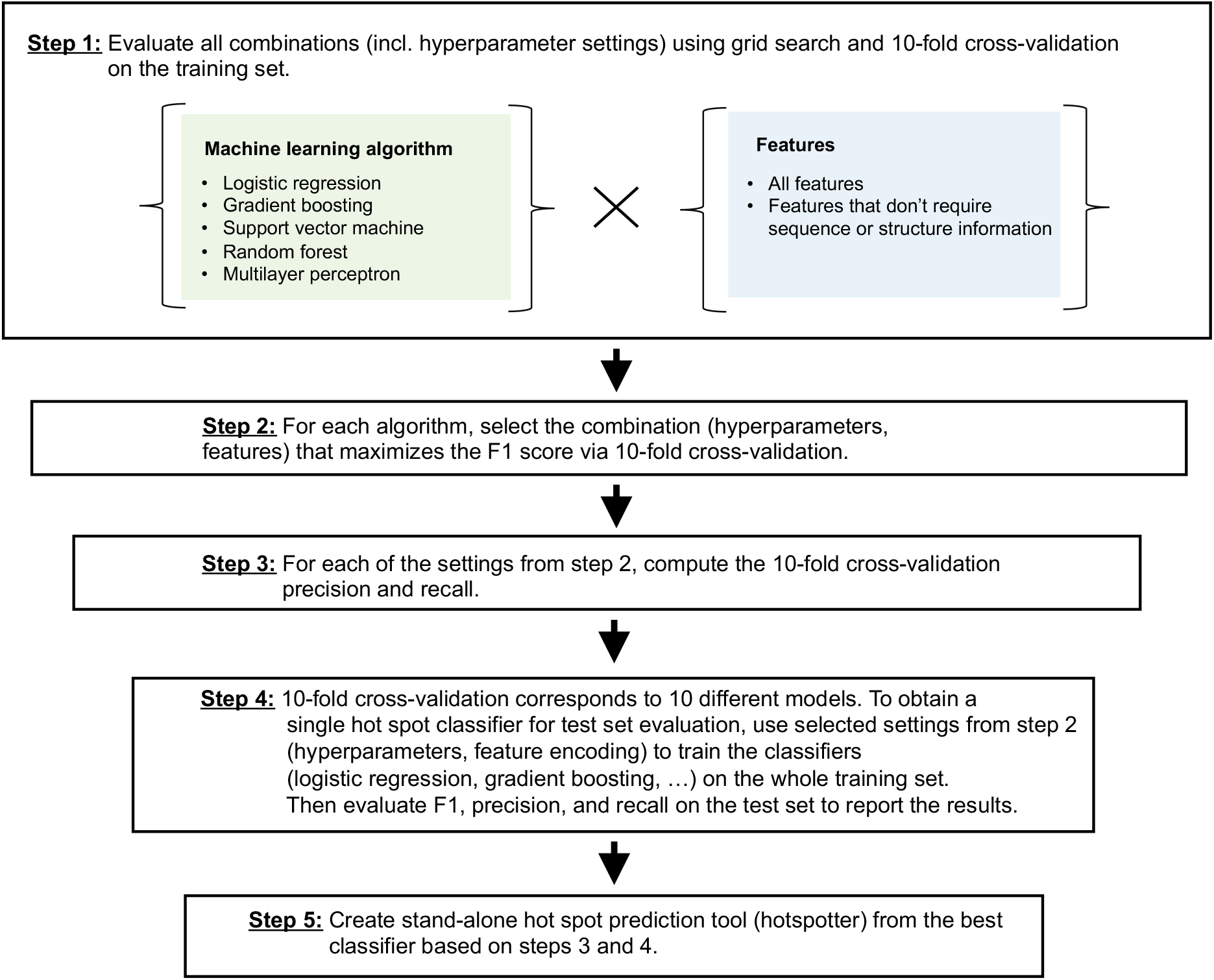
Training and validation of classifiers for *hotspotter*.

### Testing different subsets of the data to evaluate training robustness

Hyperparameters are the settings in a machine learning method that are user or default configured as fixed input to govern the learning process. Some common examples are the training set:test set split ratio and the choice of the cost or loss function to be optimized while training. We used grid search with 10-fold cross-validation to select the best choice for several other hyperparameters.

To illustrate this procedure, consider the random forest algorithm for which we evaluated several hyperparameter settings. For each hyperparameter setting, we ran 10-fold cross-validation on the training set. For a single hyperparameter evaluation run, the 10-fold cross-validation procedure partitions the training set into 10 equal-sized, non-overlapping partitions. Then, 9 partitions (90% of the training set) were used to train the random forest model, and 1 partition (10% of the training set) was used to compute the F1 score. This was repeated 10 times such that each of the 10 partitions was used for evaluation exactly once. The 10 resulting F1 scores were then averaged to obtain the score for the respective hyperparameter choice. Since the evaluation folds (90% and 10% subsets of the training data) did not overlap, 10-fold cross-validation used each data point exactly once to make full use of the data. Furthermore, since the models were fit to different subsets of the training data each time, this procedure also evaluated the robustness of the model.

After the F1 scores of all hyperparameter configurations were collected, we selected and saved the best hyperparameter configuration for each algorithm (step 2 in Figure 2). This configuration was also used to report the precision and recall values obtained via 10-fold cross-validation (step 3 in Figure 2). The code and individual settings explored for the classifiers are included in the experiments folder at https://github.com/Raschka-research-group/hotspotter. For brevity, only the best hyperparameter configurations that deviate from the default settings are listed here:

- Logistic regression: an L2 regularization penalty of C=10
- Gradient boosting: a learning rate of 0.01 and a maximum of 3 leaf nodes per tree
- Support vector machine: a linear kernel with inverse regularization parameter C=1.0
- Random forest: 1,000 tree estimators, a minimum of 2 samples per split, and a maximum tree depth of 5
- Multilayer perceptron: 2 hidden layers with 20 and 10 nodes each, tanH activation functions, and an adaptive learning rate of 0.0001 with stochastic gradient descent solver

Note that 10-fold cross-validation scores are obtained from averaging the results from the 10 individual evaluation partitions. Consequently, each 10-fold cross-validation run tests 10 different models trained on different training subsets. We re-ran each classifier on the whole training dataset using the best hyperparameter configuration to obtain a single model (final classifier) for evaluation on the independent test set, meaning the 30% of the 1,046 sites that were held back from training (step 4 in Figure 2). The code for this process is provided in: https://github.com/Raschka-research-group/hotspotter/blob/main/experiments/figure-modeling-flowchart/all-features.ipynb

### Evaluating if the sample size of training data is sufficient

To assess whether the final model obtained from step 4 (Figure 2) was trained on enough data, we constructed learning curves. A learning curve plots the classifier performance for training sets of increasing sizes. By keeping the test set constant, the learning curve shows the relationship between increasing the training sample size and the resulting classification performance on new data. For example, in Figure 3, we have plotted a learning curve for the best model (multilayer perceptron) from step 4 of Figure 2. As might be expected, the test set performance improves for larger training set sizes, though the performance asymptotes once about 60% of the entire training set has been included. This indicates that the training is thorough. The very slight positive slope of the test set performance curve (F1, measuring an equal balance of precision and recall) once 90-100% of the training set have been incorporated suggests that collecting a larger training set could slightly benefit classification. Note that the training performance of the smallest training sets (10-30% of the data) is naturally high, because it is easier for a classifier to memorize the feature-class relationship for smaller datasets. The model was overfitted to the smallest training datasets, as shown by the high training performance and low test set performance. This unfavorable characteristic can be avoided by using a larger training set. For the Results, we used all of the training data. (Note 2)

**Figure 3.**
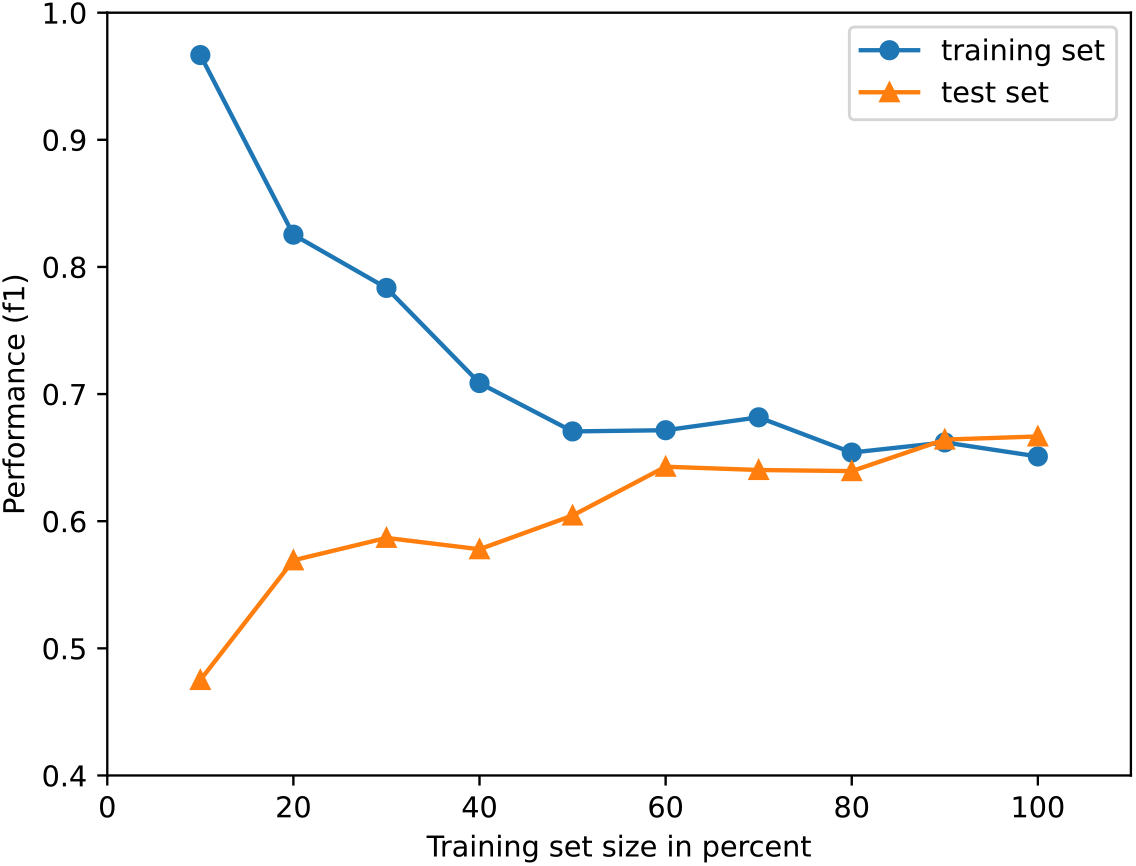
Learning curve for the all-feature multilayer perceptron classifier applied to nested subsets of increasing size from the training set. Resulting performance on the training set is graphed in blue, with results on the full test set in orange. Test set performance improvements substantially leveled once 60% of the training set had been included, indicating that the full training set sample size was sufficient.

### Applying hot spot prediction with *hotspotter* on new data

Completing the classifier optimization flowchart (Figure 2) resulted in a hot spot classifier named *hotspotter*, with a simple-to-use command line interface that researchers can apply to their own data. For protein complexes without 3D structures, a variant of *hotspotter* uses a reduced set of features as the basis for prediction: the residue type and ConSurf evolutionary conservation score for the residue, augmented by the secondary structure at this position in the protein, which can be assigned with good accuracy by AlphaFold (https://alphafold.ebi.ac.uk) or PredictProtein (https://predictprotein.org). *Hotspotter* is available at https://github.com/Raschka-research-group/hotspotter

### Choosing appropriate performance metrics to evaluate prediction quality

Accuracy, which measures the percentage of all predictions that are correct, is often used to evaluate predictors:

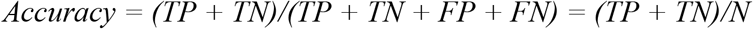

TP is the number of cases that are true positive predictions, meaning hot spots that are correctly predicted as hot spots. TN is the number of true negative predictions, meaning non-hot spots correctly predicted as such. FP is the number of false positive predictions, that is, non-hot spots incorrectly predicted as hot spots. Finally, FN is the number of false negative predictions, hot spots incorrectly predicted as non-hot spots. N is the total number of cases (TP + TN + FP + FN).

For applications like ours, in which the positive cases (hot spots) are relatively rare when the classifier is applied to new data, being buried in a large number of negative cases (non-hot spots), accuracy is actually a poor measure of predictive success. For instance, a “dummy” classifier that predicts all cases as non-hot spots while missing the cases of interest (hot spots) will obtain a high accuracy value when the number of non-hot spot sites is large. A predictor that overpredicts the number of hot spots can also attain a high accuracy score, as the non-hot spot cases will dominate the accuracy value due to their prevalence. The goal is to balance the retrieval of as many true positives (hot spots) as possible, while not overpredicting them. The statistical measures fitting this goal most closely are:

*Precision = (TP)/(TP + FP*), which measures the percentage of predicted positives (hot spots) that are actually hot spots. High precision indicates that most of the hot spots predicted are correct, corresponding to a low rate of overprediction.

*Recall, also known as sensitivity = (TP)/(TP + FN*), is the retrieval rate, measuring the percentage of known positives, i.e., hot spots, that are predicted as hot spots. High recall indicates that most of the known hot spots are predicted correctly.

F1 is the average of the precision and recall, representing an equal balance. F1 is a convenient single metric for measuring the balance between capturing the true positives while not overpredicting them, which is the fundamental goal for our classifier, *hotspotter*.

A fifth commonly used performance metric is:

*Specificity = TN/(TN+FP*), which measures the percentage of all actual negatives (non-hot spots) that are correctly predicted as negatives. Specificity is less relevant for our problem because it emphasizes correct prediction of the negative, non-hot spot case which is highly prevalent relative to hot spots in new applications to proteins. Non-hot spots (negatives) incorrectly predicted as hot spots are already evaluated in the precision metric above. Seeking high specificity in a situation where the positive cases are rare will lead to the dummy predictor issue described above: a classifier with significant under- or over-prediction of the positive category (hot spots). Here, precision, recall, F1, specificity, and accuracy were all calculated from the data using the equations above; they can also be calculated in scikit-learn. (Note 3)

### Assessing correlation between features and their relative importance for prediction

Once the features have been computed for a broad set of samples, such as the training set, it is valuable to examine the correlation between features. Including a feature that is highly correlated with another feature in the classifier does not add new information, while obscuring the interpretation of feature importance. For instance, if you include two features that are highly correlated in a linear regression classifier, a weight of 1 on one feature and a 0 weight on the other feature, or any value in-between such that the two weights equal 1, will lead to a very similar prediction. However, one feature is not much more important than the other, nor do they vary in importance; they are essentially the same feature. This phenomenon occurs to a lesser extent when two features are significantly but not highly correlated. Ideally, correlation between features is assessed early in a project, allowing the number of features to be simplified to only include the subset that are at most moderately correlated with each other. Counter to the common lore, it is typically untrue that including more features, or more detailed features, leads to better classification in protein applications, as observed later in this work. (Note 4)

The independence of two features can be tested by measuring the correlation coefficient between their values across samples in the training dataset, or across all samples in the training + test datasets, before using them in the classifier. Here, we used the Excel CORREL function to measure the linear correlation coefficient between each pair of feature columns across the set of 1,046 sites, as shown in the Results. Choice between features that are highly correlated can be made according to which of the features leads to higher precision and recall or F1 value in the classifier results.

Given a set of features used to characterize the samples for prediction and the final, trained version of the classifier, a major goal is to gain insight into which features were most important for prediction. Those features can be very useful in driving molecular design. An intuitive way to evaluate feature importance is to leave out each one, in turn, and retrain the classifier. The results before and after leaving out a feature measure the performance loss (or possibly gain) from omitting that feature. However, this approach can be time-intensive, as it requires retraining the classifier for each left-out feature. Alternatively, one can use a classifier that has already been trained, and simply shuffle or randomly reorder the data for one feature across all the test set samples, and then reapply the classifier. The prediction is then performed with the randomized values of one feature and the correctly assigned values for all the other features. This shuffling of values for the feature is repeated 5 times for robustness, and the statistics of performance across the shuffles are reported. The average performance decline from masking that feature by shuffling indicates how important the feature is. The process is then repeated for each other feature. This technique is called permutation importance, and we use the implementation described in: http://rasbt.github.io/mlxtend/user_guide/evaluate/feature_importance_permutation/ from MLxtend v0.21.0 [37]. (Note 5)

### Applying *hotspotter* to new proteins: predicting critical residues on human ACE2 for binding the SARS CoV-2 spike protein

To apply the *hotspotter* classifier to an interesting and important complex, we focused on the interaction between the human ACE2 protein (angiotensin-converting enzyme 2) and the SARS CoV-2 coronavirus spike protein. The complex between them is required for the COVID-19 virus to initiate entry into human cells. Although a detailed set of ΔΔG_binding_ values for the interfacial residues in this complex is not yet available, deep mutagenesis testing of all 20 amino acid residues in place of the wild-type residue has been performed for almost all residues between Thr 20 and Arg 518 in human ACE2 [38]. The authors rank each substitution in ACE2 by the resulting log(depletion) or log(enrichment) in binding to the SARS CoV-2 receptor binding domain, which includes the spike region that directly binds ACE2 and is used to develop COVID-19 vaccines. Enhanced binding to the CoV-2 spike protein by mutations in ACE2 is less common but possible. This is because the SARS CoV-2 and ACE2 proteins have coevolved for a limited time, resulting in interfacial contacts that are not yet optimal.

Our interest is in the ability of *hotspotter* to predict hot spots of interaction between ACE2 and the SARS CoV-2 spike, which could then be targeted by ligands to block viral entry into human cells. Thus, we focused on the high log(depletion) mutations, that is, residues where substitution by most other amino acids leads to substantially decreased binding to SARS CoV-2 (indicated in orange in Figure 1 from [38]). Similarly, non-hot spots were defined as residues in ACE2 that were tolerant of most amino acid substitutions, maintaining the ability of ACE2 to bind the spike protein (indicated in white in Figure 1 from [38]). Because we are interested in direct interactions between the two proteins, the highly-sensitive (hot spot) and highly tolerant (non-hot spot) residues in ACE2 were first filtered to identify the surface-exposed subset with relative accessibility of 10% or more in the ACE2 chain only of PDB entry 6m0j (https://rcsb.org; [20, 39, 40]. Relative accessibility was calculated in PyMOL version 2.5.2 (The PyMOL Molecular Graphics System, Warren L. DeLano and Schrödinger, LLC). The surface-accessible plus experimental high log(depletion of binding) experimental definition of hot spots in ACE2 differ from the ΔΔG_binding_ criterion used in *hotspotter* training. However, the ACE2 hot spot definitions may be more rigorous, given that they are based on more mutational data showing not only does alanine substitution significantly decrease binding, but also almost any amino acid substitution in the same position is highly deleterious.

## Results

### Assessing features for hot spot prediction

First, we analyzed the known hot spots in our dataset of 1,046 sites to ascertain whether certain residue types preferred being in hot spots relative to non-hot spots. The results in Figure 4 confirm prior work showing that tryptophan and tyrosine have a high preference for occurring in protein:protein hot spots [6, 29]. We did not find arginine to favor hot spots, as the earlier work had, while newly finding that phenylalanine has a high hot spot preference. Glycine (with no side chain) is also surprisingly common in hot spots. The top three residues (tryptophan, tyrosine, and phenylalanine) indicate the importance of bulky side chains with significant aromatic or π electron character in forming hot spots for protein:protein interaction.

**Figure 4.**
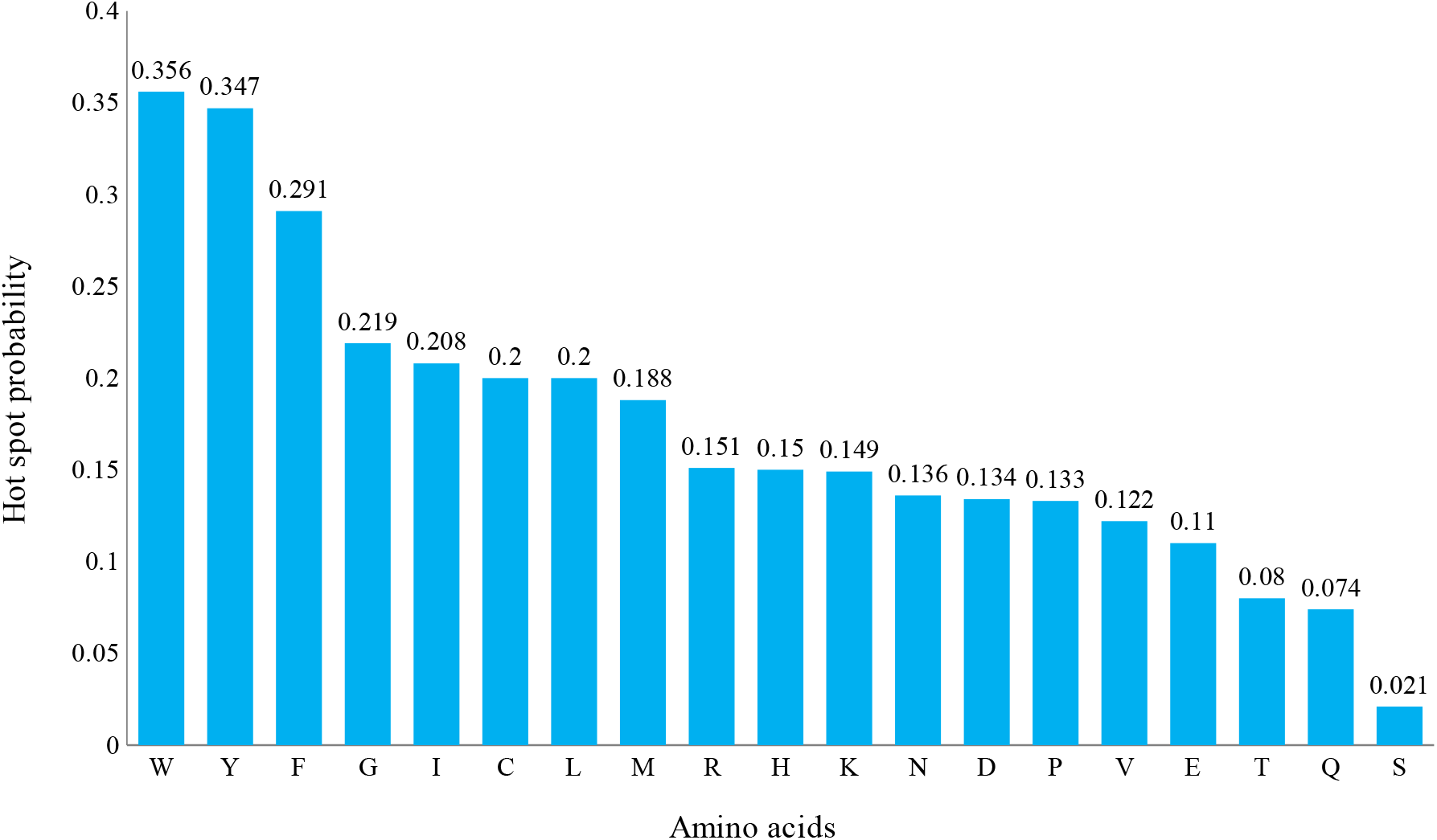
Hot spot probability for each of the nineteen common amino acids, calculated as the number of occurrences of a given amino acid with ΔΔG_binding_ ≥ 2 kcal/mol divided by the total number of occurrences of that amino acid in all the non-redundant mutations from the ASEdb, AB, BID, and SKEMPI databases. Alanine was not included, because alanine residues are not mutated in alanine scanning mutagenesis experiments.

### Evaluating classifier feature correlation in the curated mutational database

Before finalizing our feature set for predicting hot spots, we quantified the extent to which the features were correlated, by building a pairwise matrix of linear correlation coefficient values (Figure 5). The goal was to avoid incorporating in the classifier features with high correlation coefficients, approaching 1 (correlated) or −1 (anti-correlated). Such features are mostly redundant while taking effort to calculate and making it difficult to interpret relative feature importance. Ideally, all the features would have pairwise correlation coefficients near 0 (uncorrelated), indicating that each feature brings entirely new information to the prediction. The data (Figure 5) show most pairs of features have relatively low correlation (~0.3 or less). Not surprisingly, bond number, which includes the number of hydrophobic tethers (contacts) and hydrogen bonds, is moderately correlated with the hydrophobic tether count, hot spot probability, and accessible surface area (the more bonds, the more buried the residue is). Accessible surface area is moderately correlated with B’ side chain, likely because more exposed residues can be more mobile. None of these features is highly correlated with another, and so they all bring new information to the classifier. Before using these features in the training and testing of classifiers, the hot spot probability feature was replaced by the residue type in the site (without hot spot occurrence weighting), removing any possible bias from calculating hot spot probability values from our large dataset. Secondary structure type (helix, strand, turn, or other) was the eighth feature and was not included in the correlation analysis because it is non-numeric.

**Figure 5.**
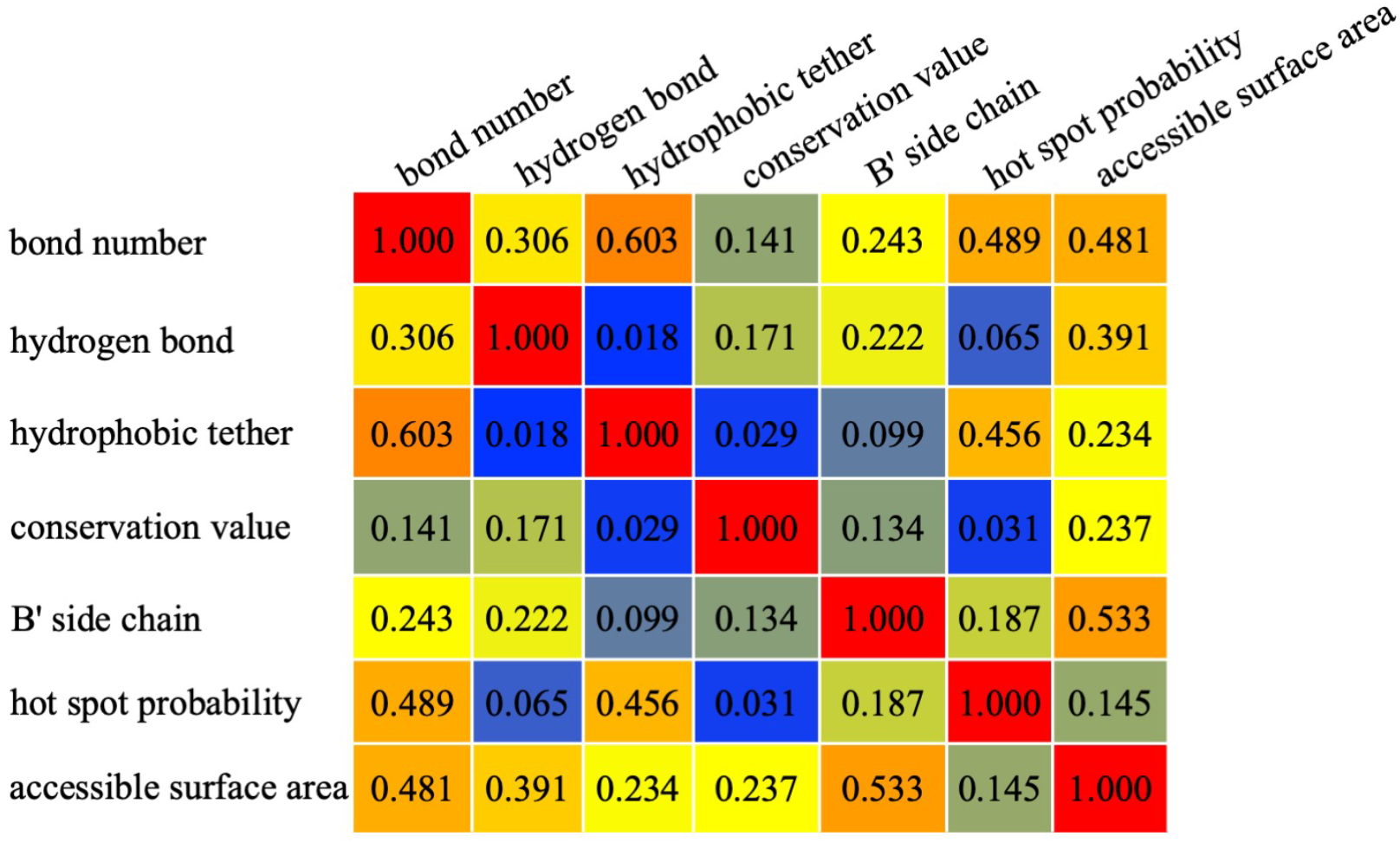
Pairwise linear correlation coefficients for the numeric features used in classification. The heatmap color scale ranges from red (high correlation) to orange (moderate correlation) to yellow (low correlation) to olive and grey (very low correlation).

### Hot spot prediction from training and testing five different classifiers on 1,046 sites

The precision, recall, and F1 results of using all eight features versus the sequence features set (residue type, ConSurf conservation score, and secondary structure type) to classify 1,046 sites as hot spots or non-hot spots were compared for five different machine learning methods (Table 2). An interesting observation in these results is that the F1 value, reflecting both precision and recall, is better (higher) for the unbiased test set than for the 10-fold cross-validation (CV) performance on the training set. This is true for all the classifiers except gradient boosting, which we noticed must be carefully tuned with fewer hyperparameters (and did) to avoid overfitting the training set. The classifiers may have been able to perform better in test set prediction due to having access to 10% more training data, relative to the 10-fold CV results which learned from 90% of the training set.

**Table 2.**
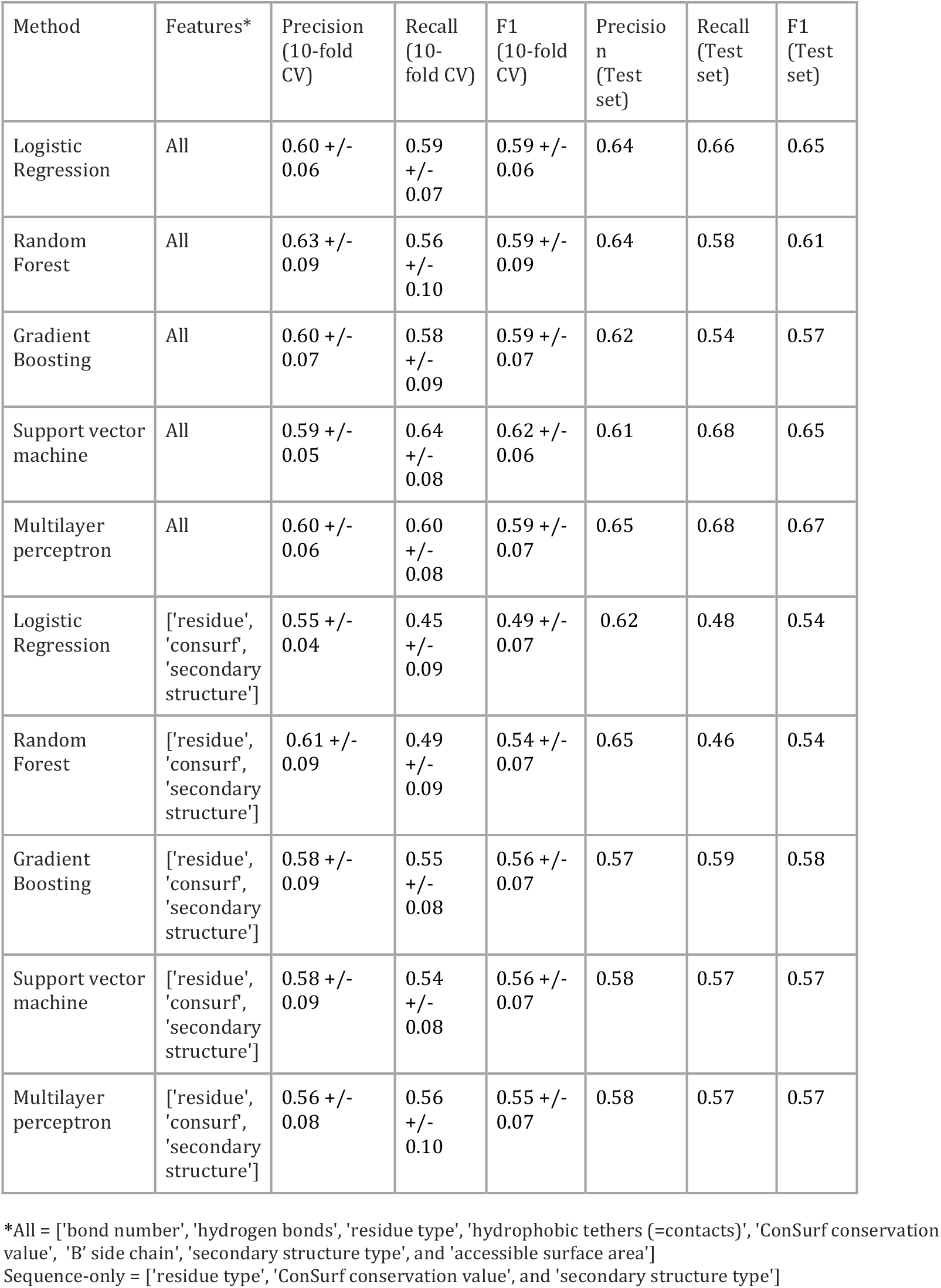
Hot spot classifier training and unbiased testing results

For all classifiers, even with non-homologous sites in the dataset randomly assigned to the training and test sets, there can be slight differences in the kinds of sites included in the training and test sets. Here, 732 sites were included in the training set and 314 in the test set, corresponding to the 70%:30% split. The large training set (which Figure 3 shows was useful) also could make its sites more difficult to predict, in that a larger dataset can include more diverse examples. Yet, the overall message from comparing the training set and test set F1 values is that the classifier results are well-balanced between training and testing, showing no sign of overfitting, which would be characterized by much better performance on the training set than on the unbiased test set. Furthermore, the classifiers trained on sequence features show almost perfect balance in performance between the training and test set, with some loss of performance on the test set relative to using all the features.

### Diagnosing if hyperparameters have been overtuned in training

To explore the phenomenon of overtuning, we evaluated 50,000 input hyperparameter configurations of gradient boosting, a classifier applied by other groups for hot spot classification. The best hyperparameter configuration resulted in an F1 score of 0.63 on the test set used in Table 2. To probe whether this hyperparameter choice was robust, we temporarily merged the training and test sets and repartitioned 70% of the merged set into a new training set, placing the remaining 30% in a new test set. The process was repeated 500 times. For each of the 500 different repartitions, the gradient boosting model was trained on the new training set and evaluated on the new test set. As shown in Figure 6, the average test set F1 performance of the gradient boosting classifier over all the repartitions was 0.46 (panel A) when a large set of hyperparameters had been optimized. However, it improved to 0.60 when the number of optimized hyperparameters was reduced to 12 (panel B). The fact that the average F1 over 500 test sets for gradient boosting with a small number of hyperparameters was 0.60, better than the F1 of 0.57 reported for the same method and hyperparameter configuration in Table 2, simply indicates that the Table 2 test set was harder than average.

**Figure 6.**
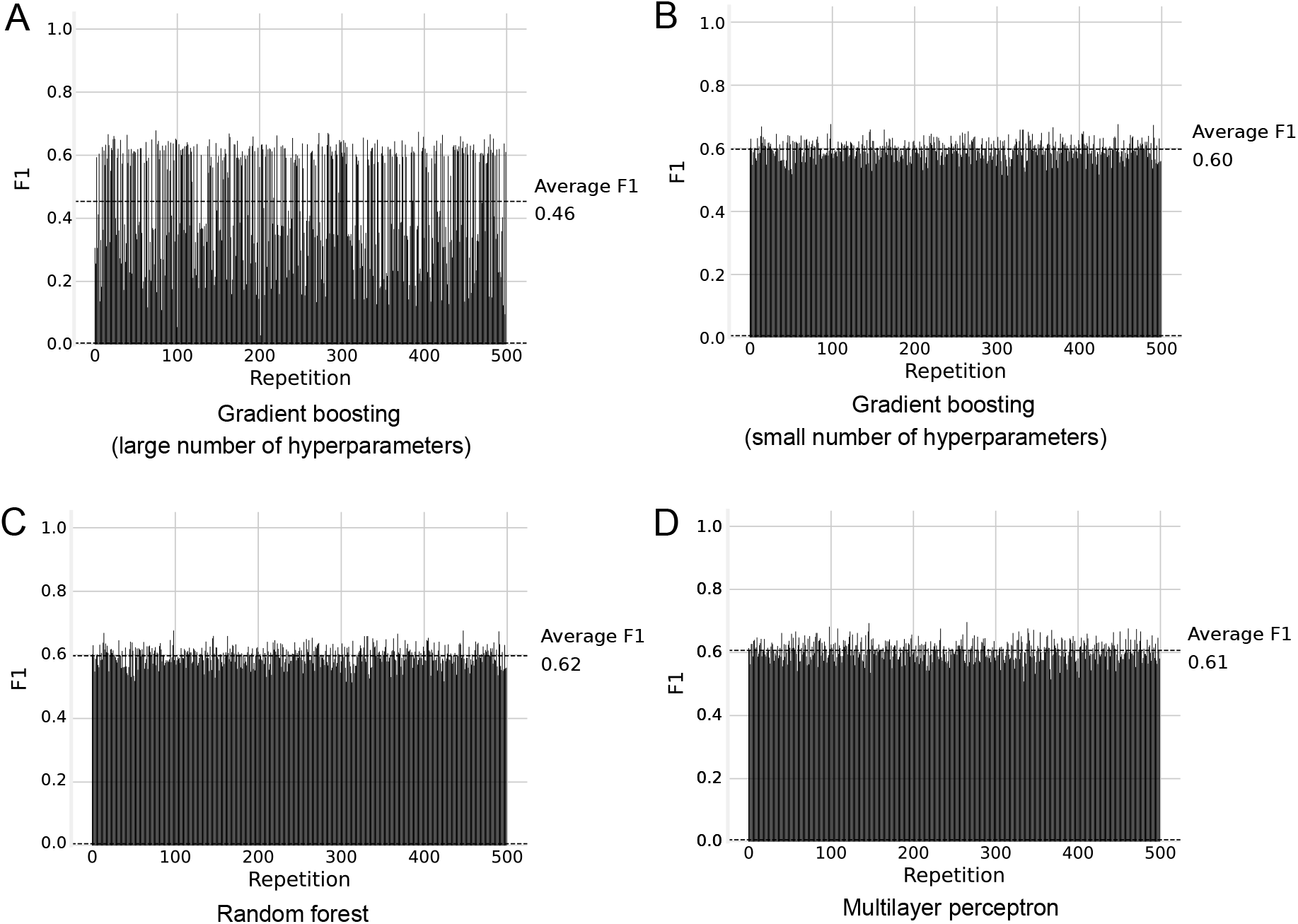
Analyzing the influence of the number of hyperparameters on classifier performance across different training and test sets. The results highlight the dangers of overfitting (often unknowingly) via tuning a large number of hyperparameters for a method on a given dataset and then applying it to new data. Each of the 500 experiments summarized along the x-axis corresponds to a new repartition of the training and test sets, as described and analyzed in the text. The data shown is for classifiers using all eight features. This analysis was repeated for the same classifiers employing the three sequence features only and showed substantially similar results.

Most importantly, when only a few hyperparameters (12) were sampled and tuned (panel B), the high variability in F1 performance across repartitions of the data (panel A), with F1 values ranging vastly between 0.2 and 0.6, was no longer encountered. For comparison, Figure 6 also shows the performance of the random forest classifier (9 hyperparameter configurations sampled) and the multilayer perceptron classifier (120 hyperparameter configurations sampled) used for the data in Table 2. Both showed high consistency in performance across the 500 runs and good average F1 scores (0.62 and 0.61, respectively). (Note 5)

Thus, 10-fold cross-validation during training is not a silver bullet against overfitting. Repartitioning the training + test dataset multiple times, as shown here, or applying the classifier to a range of new data, is invaluable for assessing whether overtuning of input parameters or overfitting to the training set has occurred. This will assess before public use whether the classifier performs consistently well on new data and is far superior to the single choice of training and test sets used in many studies, which can be biased by evaluation in the developer’s lab.

### Selecting the final *hotspotter* classifier

Considering all the results, the multilayer perceptron (MLP) proved to be the most robust of the five classifier types, showing good balance between precision and recall for the 10-fold crossvalidation on the training set and also on the unbiased test set, similarly high F1 values between the training and test sets, and low variance in F1across the 500 repartitions of the training and test sets. Figure 7 summarizes the true positive (hot spot), false positive, false negative and true negative (non-hot spot) predictions of the MLP classifier across the 732 samples in the training set and 314 samples in the test set, for all-feature training (top row) and for sequence features only (bottom row). The number of TP hot spot and TN non-hotspot predictions are significantly greater for the sequence feature version than for the all-feature version on the training set but worse on the test set – particularly in the loss of 15 true hot spots.

**Figure 7.**
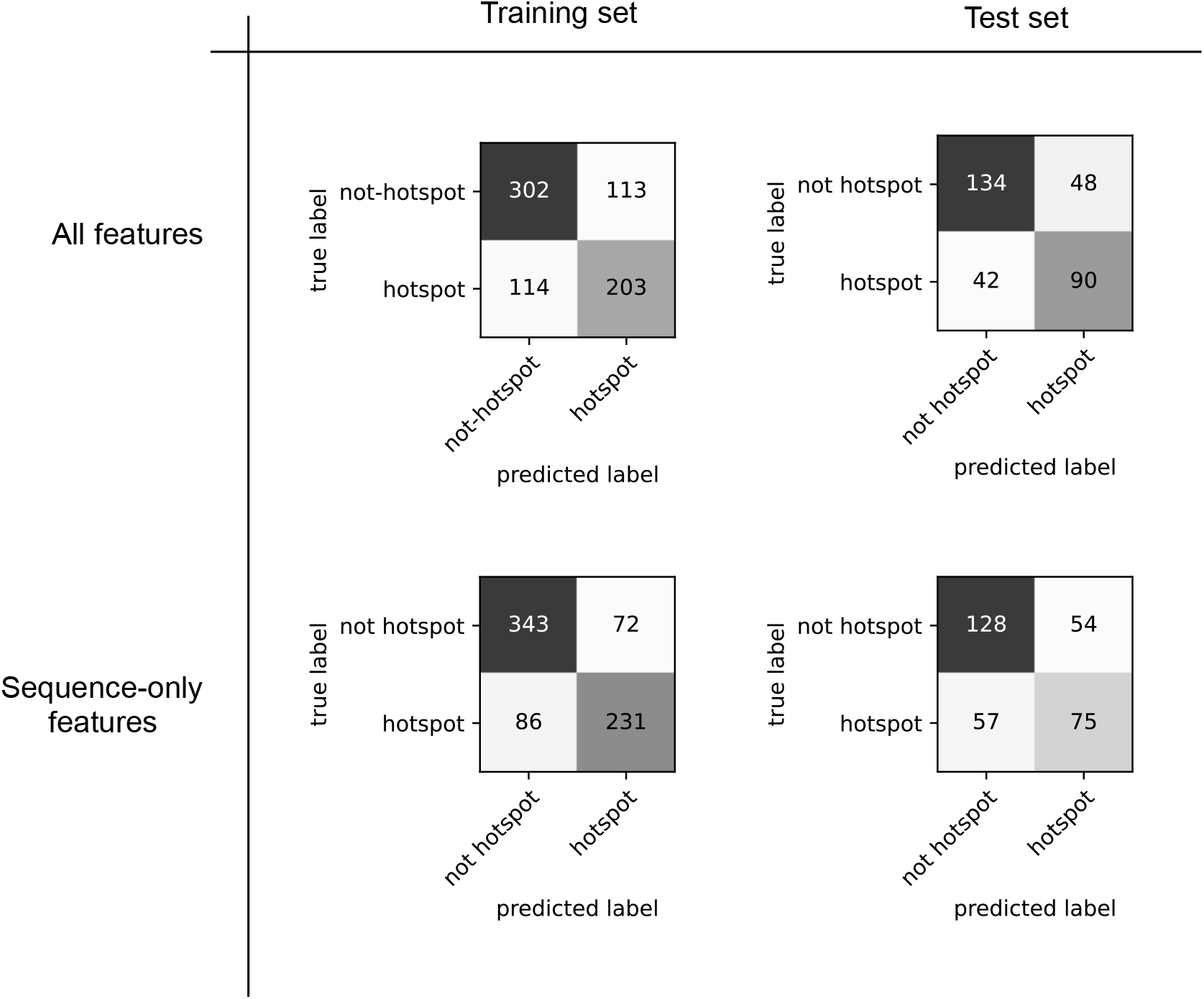
Confusion matrices listing the four categories of results from MLP classifier training (left column) and testing (right column). Within each matrix, the top row lists the number of true negative predictions (non-hot spot residues predicted as non-hot spots) followed by the false positives (non-hot spot residues predicted as hot spots), and the bottom row lists the false negatives (hot spot residues predicted as non-hot spots) followed by the true positives (hot spot residues predicted as hot spots). The first row of classifier results is based on all eight features, and the second row is based on the three sequence features. These data are interpreted in the section, Selecting the final *hotspotter* classifier.

When the entire dataset of 1,046 sites was evaluated by summing the training and test statistics for each of the categories (TN, FP, FN, and TP) in Figure 7, overall the performance of the multilayer perceptron using sequence features seemed superior, correctly predicting 471 non-hot spots (TN) and 306 hot spots (TP), relative to 436 correct non-hot spots and 293 correct hot spots for the all-feature classifier. However, 500-fold repartitioning of the 1,046 sites into new training and test sets followed by retraining the classifier for each new training set showed the all-features classifier to have consistently higher performance. All-features classification had an average F1 value of 0.60 across the 500 runs, whereas the average F1 from sequence features classification was 0.54. Both showed low variability in F1 (< 0.05) across the 500 runs, which is crucial. The all-features classifer correctly predicted 293 of the hot spots out of 454 in the database. The all-features and sequence feature versions of *hotspotter* are both available at https://github.com/Raschka-research-group/hotspotter (Note 6)

### Defining site features most important for hot spot identification

From using the shuffling method to quantify feature importance for the all-feature MLP classifier, accessible surface area and the ConSurf measure of evolutionary conservation were found to be the most important features for distinguishing hot spots from non-hot spots (Figure 8). The sequence feature MLP classifier uses ConSurf evolutionary conservation, secondary structure type, and residue type to predict hot spots; the other features were not included because they require detailed 3D structural information.

**Figure 8.**
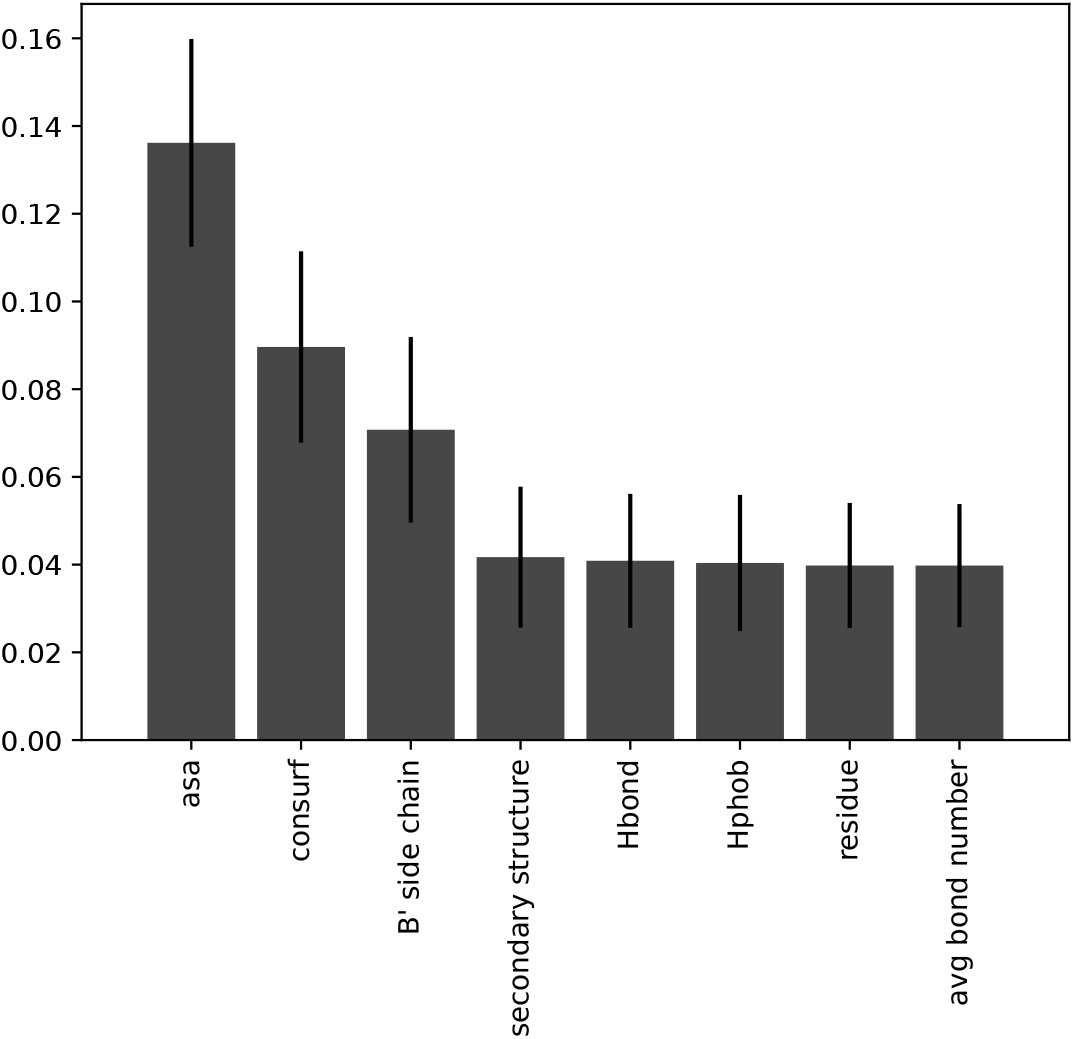
The relative importance determined by permutation importance of eight different features for the prediction of protein:protein hot spots and non-hot spots by the multilayer perceptron *hotspotter*. One feature’s values in the training set are randomly shuffled, followed by classifying hot spots or non-hotspots using the shuffled feature and the correct values for all other features, and then the F1 value is measured. The average and standard deviation in F1 loss upon 5 rounds of shuffling and classification are reported on the y-axis, with higher values indicating more important features for classification.

### *Hotspotter* prediction of hot spots on human ACE2 receptor for SARS CoV-2 spike binding

*Hotspotter* was applied prospectively to a case different from those used in training, both in the type of interface and the way the hot spots are defined experimentally. The relatively small, crescent-shaped interface with the SARS CoV-2 spike protein centers on the curved helix formed by residues 20-53 in ACE2, which is crucial for virus binding and infection of human cells. Known hot spots were experimentally defined [38] as ACE2 residues that could not be substituted without substantially abrogating SARS CoV-2 spike binding (as discussed in the Methods here) while also having at least 10% relative surface accessibility in the 2.45 Å resolution structure of the spike-bound conformation of ACE2 (PDB entry 6m0j; [39]). Following these criteria, there are eleven experimentally defined hot spot residues in ACE2: Thr 20, Ile 21, Phe 28 (especially sensitive to mutation), Glu 37, Asp 38, Gly 326 (especially sensitive), Trp 349, Asp 350, Lys 353 (especially sensitive), Gly 354 (especially sensitive), and Phe 390.

Results of *hotspotter* prediction in ACE2 are shown in the context of its complex with spike (Figure 9) and summarized here:

- The six correctly predicted hot spots (true positives) are shown in magenta in Figure 9. *Hotspotter* identified the critical hot spot triad of Glu 37, Asp 38, and Lys 353 side chains forming a cluster of direct interactions with the spike protein at the center of the interface as well as two of the four sites that were especially sensitive to mutation. The additional three predicted hot spots – Phe 28, Trp 349, and Asp 350 – are situated one layer away from the interface, in or near the ACE2 helix that binds to the CoV-2 spike protein.
- The five experimentally defined hot spots that were missed are shown in gray. Two are glycine residues (Gly 326 and Gly 354) in direct contact with the spike protein, while the other three are not near spike (Thr 20, Ile 21, and Phe 390). The intolerance of mutation with respect to spike binding for Gly 326 and Gly 354 likely reflects that side chains added in these positions would sterically hinder forming a complex with spike and also reduce the flexibility of the ACE2 main chain, potentially constraining the conformations of these loops needed to bind spike. Mispredicting the two glycine residues in direct contact with spike suggests that specific training to distinguish between Gly non-hot spots and interfacial Gly hot spots, which comprise a significant percentage of all glycine residues in the ΔΔG_binding_ dataset, could potentially benefit from additional features like main-chain flexibility.

**Figure 9.**
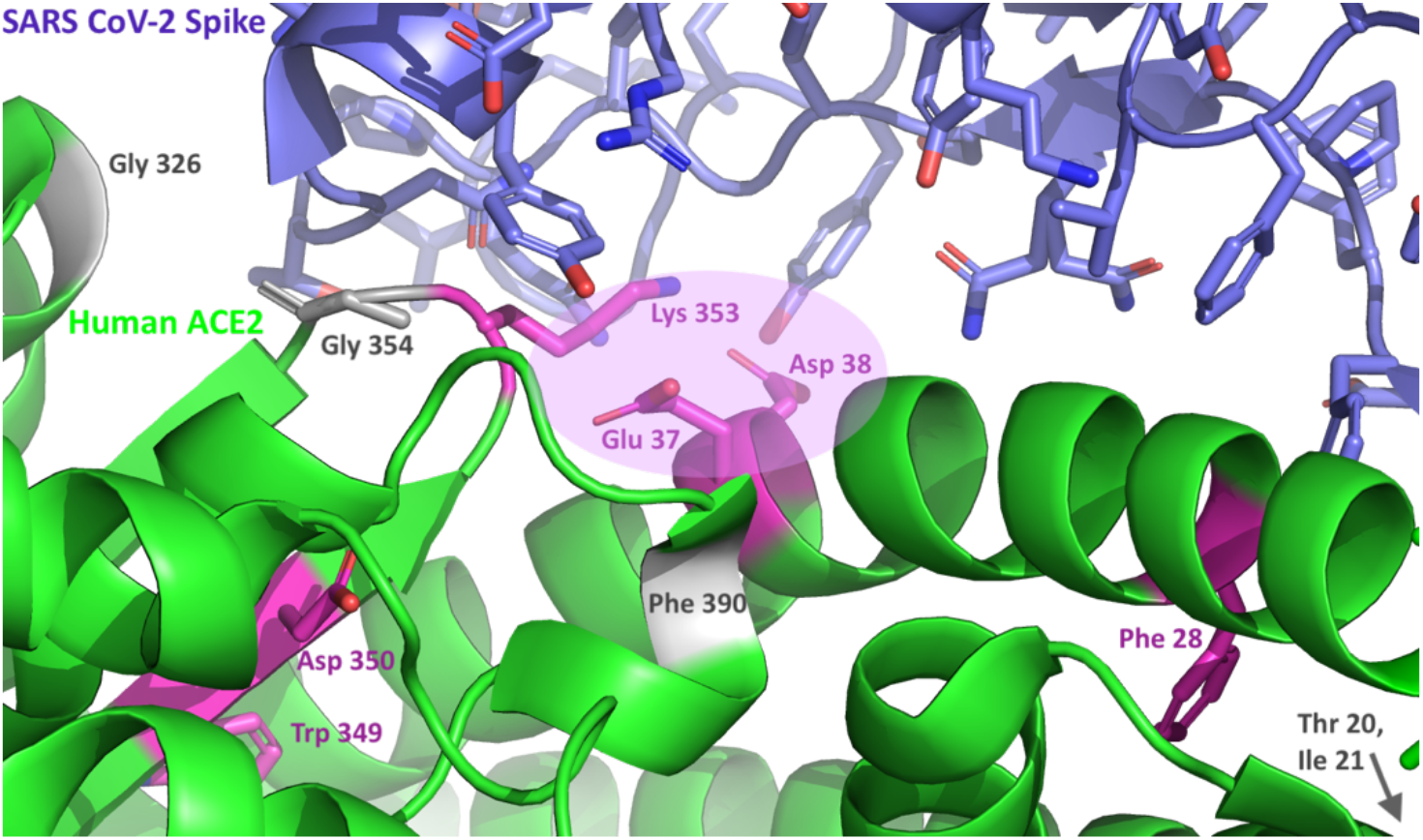
*Hotspotter* prediction of interaction hot spots on the human ACE2 protein interface with the SARS CoV-2 spike protein, which promotes COVID-19 infection. ACE2 is shown in green ribbons (lower half of the figure), and the SARS CoV-2 spike protein appears in blue ribbons (at top), from PDB entry 6m0j, a 2.45 Å resolution crystal structure of the complex [39]. The six true positive predictions of hot spot residues are shown in magenta (three interfacial and three non-interfacial), including the critical triad of Glu 37, Asp 38, and Lys 353 side chains that form a cluster of direct interactions with the CoV-2 spike protein at the center of the interface. Missed hot spot residues, defined based on mutagenesis only and not surface or interfacial exposure, are shown in gray; two are Gly residues (326 and 354) that are interfacial with spike, while the other three (Thr 20, Ile 21, and Phe 390) are a layer away from the binding site. These results point to the importance of screening hot spot predictions for residues actually occurring on the surface or in the interface. Ideally the mutagenesis/binding energy database used to train the classifier will be filtered first to only include surface or interfacial residues, allowing better training on the features of *interfacial hot spots* instead of residue changes that destabilize the protein.

### Focusing hot spot databases on surface and interfacial residues to improve their usefulness

The list of eleven experimentally defined ACE2 hot spots, even when filtered for surface exposure, includes three residues (Thr 20, Ile 21 and Phe 390) that are not close to ACE2’s protein partner, CoV-2 spike. Such residues distal from interfaces are seldom of interest for protein or ligand design because their mutations likely destabilize the protein or its conformation in unpredictable ways, rather than strategically making direct interactions with a protein partner. They would be considered structural integrity hot spots rather than protein interaction hot spots. (Note 7)

The list of experimentally defined hot spot residues in ACE2 and the proteins in mutational and ΔΔG_binding_ databases is far greater than the actual number of interfacial hot spot residues, because most of the ΔΔG_binding_ databases do not take advantage of structural information available from the PDB or AlphaFold predictions to focus on surface accessible residues, better yet, interfacial residues, as hot spots. Interfacial residues still vary in ΔΔG_binding_ upon mutagenesis, and the ultimate goal is to distinguish the most energetically important interfacial residues from other interfacial residues.

Thus, a valuable way forward for improving hot spot prediction in general will be to filter the ΔΔG_binding_ databases down to residues that are surface-exposed and interfacial. This will be invaluable in solving a current problem throughout the field: by optimizing the prediction of hot spot residues to match the ΔΔG_binding_ values in datasets that mostly do not distinguish between buried, surface, and interfacial residues, predictors have been trained to identify so-called hot spots that are actually buried in the protein or far from the binding site, as well as the typically small number of true interfacial hot spot residues. Filtering for surface or interfacial context of residues can be done following prediction, as we have done. This improves results but does not undo the bias in training (based on instances including in the training set) to identify many “hot spots” that are far from the surface or molecular interface.

### Lessons learned for training and test set design for improving hot spot prediction

To correctly define hot spots experimentally, structural destabilization of the mutated protein first needs to be ruled out. While performing protein folding and stability experiments for each mutation is not feasible computationally, we know that well-exposed sites on individual protein structures tend to be tolerant of mutations. Buried sites do not participate in direct protein:protein contacts and also tend to be intolerant of mutations, particularly by residues with different chemical character or larger size than the wild-type residue. Thus, a reasonable way to rule out mutations likely to destabilize the protein is to rule out buried residues included in experimental lists of hot spots, and instead focus on the subset of residues that contribute significantly to the solvent-accessible surface of the protein and are available to bind the protein partner.

Making the distinction between buried sites and solvent-exposed sites in proteins without experimental 3D atomic-resolution structures is now much easier, due to recent breakthroughs by the 3D structural modeling software AlphaFold, which has similar accuracy to experimental structure determination in a majority of cases (https://alphafold.ebi.ac.uk; [41]). AlphaFold can be used to predict the atomic structure of a protein, and that structural prediction can then be used to define which residues are substantially buried and likely to affect structural stability if mutated, versus which residues are surface-exposed enough to directly interact with another protein or ligand. An easy to apply and robust criterion for surface exposure is to identify those residues contributing at least 5 Å^2^ (roughly 1.5 carbon-carbon bonds in length and width) to the *solvent accessible surface area* (SASA, not ASA or relative surface area) of the protein *in the absence of its binding partner*. The solvent-accessible surface area (SASA) of a protein is like a shrink-wrapped version of a protein, in which each residue’s contribution to the surface is measured by its SASA area. SASA turns out to be a better criterion for surface accessibility than the relative surface area calculated on a per-residue basis, which we now know does not fully measure the effect of neighboring residues in burying a residue. SASA can be used to filter a list of experimentally-defined potential hot spot mutations to include only the substantially surface-exposed sites for training, testing and new-case predictions of hot spots by classifiers. This is likely to significantly reduce the currently misleading training of classifiers on buried sites labeled as “hot spots”, which then leads to classifiers’ incorrect identification of buried residues as protein:protein interaction hot spots when applied to new proteins. Focusing on surface sites will also greatly reduce the number of mutagenesis experiments and result in more stable mutant proteins.

## Conclusions

A major goal of the research presented here is to show the steps of validation needed to develop a robust predictive tool (in this case, a machine learning classifier) that is very likely to perform similarly well to the published results when applied by others to new cases. Dazzlingly good results on a single dataset can be achieved by extensive tuning of a classifier by an expert lab, but the results here show that very process often leads to incomplete learning of the patterns of characteristic features of positive and negative cases, and poor performance on new data.

The ways of evaluating predictive methods shown here are equally useful for evaluating publications on existing methods for their robustness in data curation, choice of performance metrics, and thoroughness in unbiased testing on a variety of data. Finally, we show that a successful classifier for a very hard problem – identifying the few most energetically important interaction residues for one protein to bind to another – can be identified from a large, noisy database by using available machine learning methods and a small number of intuitive features. Part of the process is identifying which features are most important for identifying key binding residues, which advances not only the method development but also the designability of protein binding sites and our ability to block them.

Along with the resulting ability of the *hotspotter* classifier to detect two-thirds of the experimentally defined hot spots in 97 protein complexes, analysis of the missed and overpredicted cases shows that in the future, training the classifier on only surface-exposed residues has the potential to greatly improve true protein:protein interaction hot spot prediction. This is because the experimental hot spot databases surprisingly include many buried residues unavailable to bind another protein. The code for experiments performed in this work, the feature data used for training and testing, and the resulting hot spot prediction classifiers are available for public use on GitHub (https://github.com/Raschka-research-group/hotspotter).

### Notes

A summary of key points based on our experience:

1. Curate the data to remove redundancy within the training or testing set and between training and testing sets. Make sure your data from different sources are on the same scale and can be pooled. Using l arge, diverse datasets for both training and testing will lead to greater reliability in new applications. Remove buried sites from consideration, by including only residues with SASA of at least 5 Å^2^.
2. Validate classifier(s) on increasing subsets of the training and test sets and determine a learning curve to assess whether enough training data has been used to optimize training performance, and also whether performance on the test set would benefit from more training samples.
3. Select performance metrics that ensure your classifier is solving the problem you intend to solve. If the cases you wish to predict are relatively rare compared to the number of cases being tested (for instance, interaction hot spot residues or residues important for catalytic function), then your metric for success should emphasis the capture of the active cases and penalize overpredicting them. True positives are important while minimizing false positives, which can quickly outnumber the true positives when the test set is large. In such a case of rare true positives, a “dummy classifier” that predicts all cases as inactive (true negatives and false negatives) will apparently perform very well on the dataset if you use accuracy (total number of correct predictions, or TP plus TN, divided by the total number of cases), while not learning the information needed to capture the rare TP cases. F1, an equal balance of precision and recall, is a good metric for rewarding detection and avoiding overprediction when a small number of true positives are buried in a large dataset.
4. Use only the subset of features that is discriminatory (improves prediction) for classification. Using more features, especially features that are irrelevant to the problem or are significantly correlated with each other, introduces noise into the prediction and an inability to assess the relative importance of features.
5. Test the usefulness or information content of each feature by leaving out each feature or shuffling the values for a given feature prior to classification. Too many features can cause overfitting to the training data by increasing the number of fittable features (parameters) relative to the sample size of the data. This, along with using too many hyperparameters (preset before running a classifier) will result in a classifier that performs poorly on new data. This can be avoided by testing a given classifier configuration across many repartitions of the pooled training and test sets to measure their quality and consistency of performance.
6. Share your data and methodology with others, and they will return the favor. This advances the field and enables comparative analyses on the same data. You will learn from feedback on whether your method is easy for a non-expert to use, and how well it works on a much broader range of problems than you can test on your own.
7. Check your results to see if they match your knowledge on specific cases. This gives new insights into the biology or underlying mechanism of the process and provides the basis for effectively designing new molecules with desired functions (e.g., strengthening binding or preventing binding without unfolding the protein). If the importance values of different features are counterintuitive – for instance, if buried residues appear to be more important than surface residues as hot spots of protein:protein interaction in your prediction – this can point to problems in the classifier itself. Such problems can include: using inappropriate data for training, relative to the problem you want to solve; training to predict data that has an opposite sign from what you thought (e.g., higher ΔΔG_binding_ means worse binding, not better binding); incorrectly using the software, which you can test by using sample data with known results from the developer; or mis-indexing of the input data, such that the wrong features are associated with residue numbers in the training or test data.

A sanity check of all results, that is, comparing them to ensure the trends are what an expert already knows (such as interaction hot spots being at interfaces, rather than buried inside a protein), is crucial. This also gives an advantage to classifiers that provide a clear model of which features are important (rather than being black-box-like). Another way to ensure the method you are using is a good choice is to use it on very well characterized cases where you already know the answers, to assess the quality of performance. Ideally that problem is related to the problem you really want to solve, providing more confidence that the method is appropriate.

## Acknowledgments

We sincerely thank Prof. Jerry Tsai for providing a copy of his Binding Interface Database.

